# Isolation and characterization of bacteriophages against IMP-6-producing *Klebsiella pneumoniae* isolated from clinical settings in Japan

**DOI:** 10.1101/2022.11.05.515272

**Authors:** Kohei Kondo, Satoshi Nakano, Junzo Hisatsune, Yo Sugawara, Michiyo Kataoka, Shizuo Kayama, Motoyuki Sugai, Mitsuoki Kawano

## Abstract

Carbapenemase-producing *Enterobacteriaceae* (CPE) are one of the most detrimental species of antibiotic-resistant bacteria worldwide. Phage therapy has emerged as an effective strategy for the treatment of infections caused by CPE pathogens. In west Japan, the increasing occurrence of *Klebsiella pneumoniae* harboring the pKPI-6 plasmid, which encodes *bla*_IMP-6_, is a growing concern. To manage such major antimicrobial-resistant pathogens, we isolated 29 novel phages from sewage in Japan, targeting 31 strains of *K. pneumoniae* and one strain of *Escherichia coli* harboring the pKPI-6 plasmid. Electron microscopy analysis indicated that of the 29 isolated phages, 21 (72.4%), 5 (17.2%), and 3 (10.3%) belonged to *Myoviridae, Siphoviridae*, and *Podoviridae*, respectively. Host range analysis revealed that 20 *Myoviridae* members in isolated phages infected 25–26 strains of *K. pneumoniae*, indicating that most of the isolated phages have a broad host range. The *K. pneumoniae* Kp21 can only be infected by phage øKp_21, while Kp22 can be infected by more than 20 phages. We applied a phage cocktail, which consists of 10 phages, against Kp21 and Kp22 and found that the phage cocktail delayed the emergence of phage-resistant bacteria for Kp21 strain but not for the Kp22 strain. Furthermore, phage-resistant Kp21 (Kp21r) became prone to be infected from other bacteriophages as a “trade-off” of resistance to phage øKp_21. Our proposed phage set has an adequate number of phages to combat the *K. pneumoniae* strain isolated in Japan. Notably, our work demonstrates how a suitable phage cocktail diminishes the occurrence of phage-resistant bacteria.

**Importance:** *Klebsiella pneumoniae* harboring the plasmid carrying *bla*_IMP-6_ is becoming an increasingly hazardous species in Japan. We collected and characterized 29 novel bacteriophages that infect *K. pneumoniae* carrying the pKPI-6 plasmid, isolated in clinical settings of west Japan. Our phages showed broad host ranges. We applied a phage cocktail treatment constructed from 10 phages against two host strains, Kp21 and Kp22, which show different phage susceptibility patterns each other. Although the phage cocktail delayed phage-resistant Kp21 emergence, the emergence of phage-resistant Kp22 could not be delayed. Moreover, phage-resistant Kp21 became sensitive to other phages, which did not originally infect wild-type Kp21. Our study demonstrates how a suitable phage cocktail can diminish the occurrence of phage-resistant bacteria.

## Introduction

Carbapenemase-producing *Enterobacteriaceae* (CPE) are high-risk bacteria in clinical settings globally. *Klebsiella pneumoniae*, a member of the family *Enterobacteriaceae*, causes nosocomial infections and is one of the most common causes of life-threatening infections caused by multidrug-resistant bacteria worldwide (1). *bla*_IMP_ genes are classified as class B metallo-β-lactamases. *bla*_IMP-1_ and *bla*_IMP-6_ genes are predominantly detected in CPE isolated from Japan (2, 3), whereas other types of carbapenemases (NDM, KPC, and OXA-48) are mainly detected in CPE isolated from other countries (4). *Klebsiella pneumoniae* with the pKPI-6 plasmid encoding *bla*_IMP-6_ (5), which is susceptible to imipenem but resistant to meropenem, has become increasingly common in clinical settings in west Japan since their emergence in 2009 (6). These strains are therefore of major concern in clinical settings because of the inappropriate selection of antibiotics for treatment.

Recently, research on the use of bacteriophages as an alternative for the treatment of infections caused by antimicrobial-resistant bacteria has become increasingly prevalent (7). Phage therapy targeting *Staphylococcus aureus* (8) and *Mycobacterium tuberculosis* (9) has been administered successfully to patients. Furthermore, a recent study has demonstrated effective phage therapy targeting CPE in clinical settings (10). Thus, phage therapy is now recognized as a highly reliable strategy to combat nosocomial pathogens.

Following the use of phages against bacteria, phage-resistant bacteria have emerged (11) *in vitro* (12) and *in vivo* (13). The phage cocktail strategy, which consists of several types of phages, is often used to prevent the emergence of phage-resistant bacteria. A phage bank is useful to quickly apply the phage cocktail in clinical settings (14), especially in emergency cases. As national phage banks are pertinent for the instant management of contingent nosocomial pathogen outbreaks, several countries have constructed public phage banks for the efficient use of phage therapy (14, 15). However, there is no public phage bank optimized for the trend of antibiotic-resistant bacteria in Japan.

In this study, we isolated and characterized 29 bacteriophages targeting IMP-6-producing *K. pneumoniae* and *Escherichia coli* clinical isolates as the first step in constructing a public phage library in Japan. We also describe the mechanisms by which phage cocktails reduce the emergence of phage-resistant *K. pneumoniae*.

## Results

### 1. Phage hunting and morphological analysis of novel bacteriophages

We performed phage hunting from the sewage system in west Japan, which yielded 29 phages against 32 *K. pneumoniae* and one against *E. coli* (Ec1) isolates harboring pKPI-6. Each phage name number indicates the corresponding host number. For instance, øEc_1 and øKp_1 phages were isolated from *E. coli* Ec1 and *K. pneumoniae* Kp1, respectively, as their corresponding hosts. All phage-corresponding hosts combinations are listed in Table S1. We did not discover appropriate phages against *K. pneumoniae* (Kp2, Kp6, Kp25, Kp28, and Kp29). Morphological analysis using electron microscopy indicated that 21/29 (72.4 %) isolated phages belonged to *Myoviridae*, 5/29 (17.2 %) belonged to *Siphoviridae*, and 3/29 (10.3 %) belonged to *Podoviridae* (Fig. 1). All TEM images of *Myoviridae* were shown in Fig. S1 The transmission electron microscopy image strongly suggested that øEc_1 belonged to *Podoviridae* and C3 morphotype (honeycomb-like) phages (16–18), which possess an elongated head (height, 136.6 nm ± 1.8 nm; width, 61.7 nm ± 3.6 nm; tail, 15.8 nm ± 2.3 nm) (Fig. 1). øEc_1 formed turbid plaques on the *E. coli* Ec1 strain. øKp_21 was classified as *Myoviridae* (height, 133.8 nm ± 3.1 nm; width, 137.1 nm ± 1.1 nm; tail, 109.7 nm ± 1.0 nm) (Fig. 1). Furthermore, øKp_21 had a branched tail (tail spike) fiber and formed clear plaques on the lawn of *K. pneumoniae* Kp21.

**Fig. 1.**
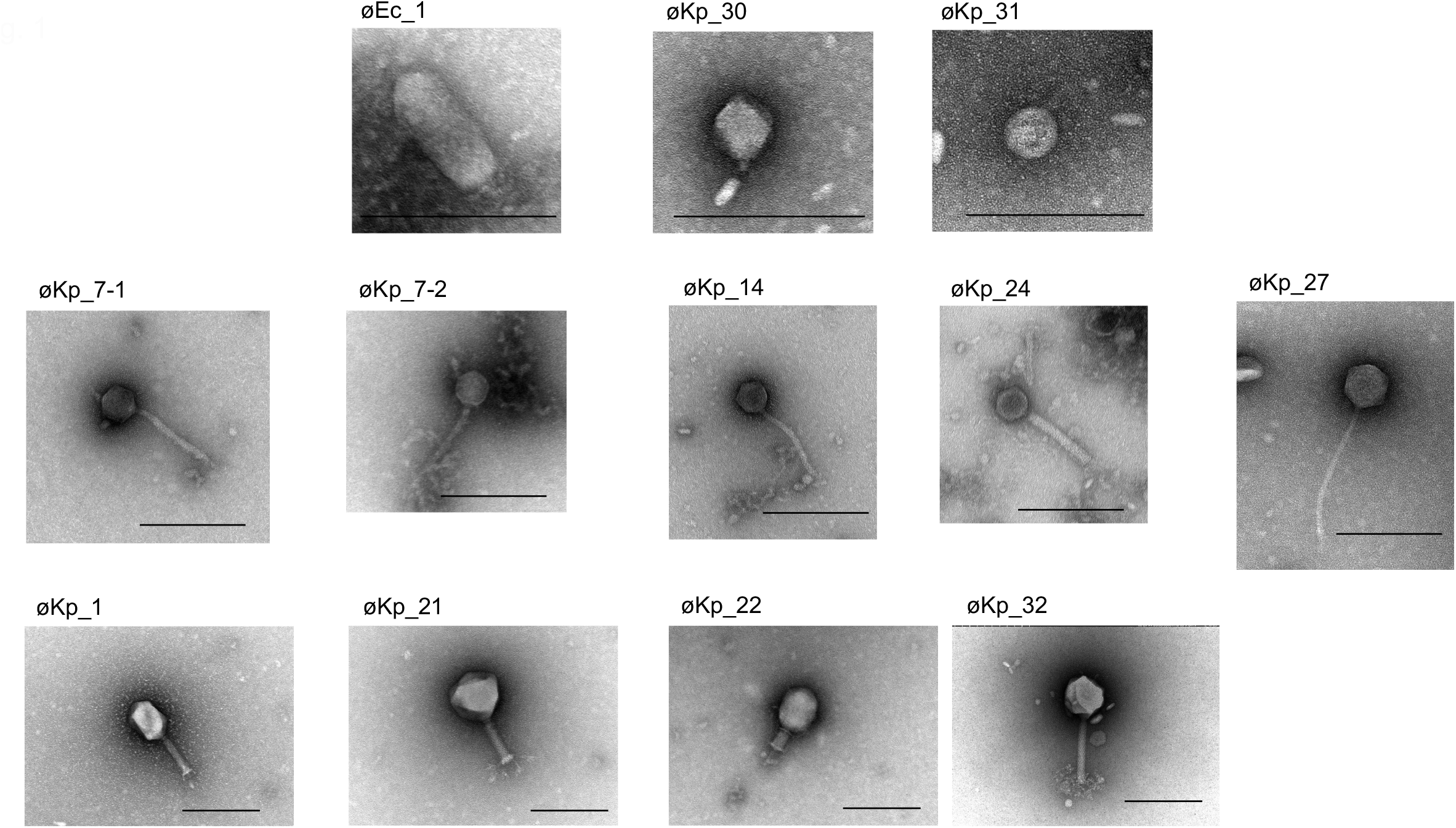
Transmission electron microscopy images of 29 isolated phages. Each sample was negatively stained and magnified at ×50,000. øKp_1 and øKp_22 are shown as representative *Tevenviridae*. All *Podoviridae* and *Siphoviridae* are shown. The bar represents 200 nm in individual images.

### 2. Host range determination and analysis of the correlation between plaque size and efficiency of plating (EOP)

We next determined the host range and its EOP for all phage–host combinations (Fig. 2) (Table S2). We selected two standard strains (ATCC BAA 1705, ATCC BAA 1706) as the control for *K. pneumoniae*. Although no novel phages against five hosts (Kp2, Kp6, Kp25, Kp28, and Kp29) were isolated, the host range experiment indicated that several isolated phages formed plaques against Kp2, Kp6, Kp25, and Kp29 but not Kp28 (Fig. 2). Phages øKp_8 and øKp_17 showed the broadest host range (26/32 *K. pneumoniae* host strains). Most phages (øKp _3, øKp _5, øKp _9, øKp _10, øKp _12, øKp _13, øKp _15, øKp _16, øKp _18, øKp _19, øKp _20, and øKp _26) showed plaque formation on 25 host *K. pneumoniae* strains, indicating that these phages have a broad host range (infecting ≧ 25 host strains). In contrast, several phages showed a narrow host range (infecting ≦ 4 host strains). For example, øKp_31 infects Kp15, Kp17, Kp26, and Kp31. øKp_27 (*Siphoviridae*), øKp_30 (*Podoviridae*), and øKp_32 (*Myoviridae*) infect only Kp27, Kp30, and Kp32, respectively (Fig. 2). Overall, our phage set can handle the *K. pneumoniae* isolated in clinical settings and has an adequate number of phage types to construct a phage cocktail against *K. pneumoniae* in this study.

**Fig. 2.**
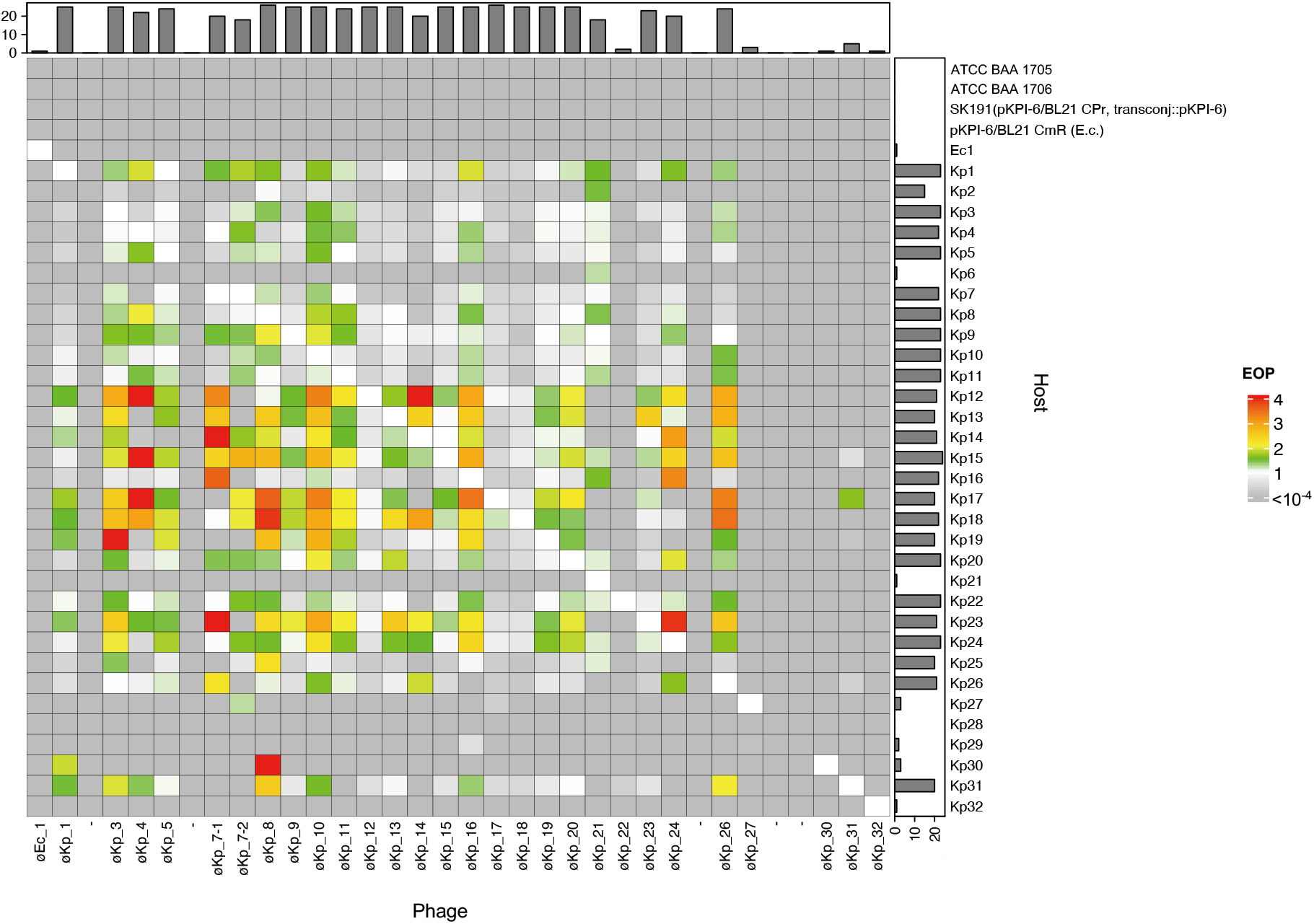
Heatmap of the host range in each phage. X and Y axes represent phages and host strains, respectively. *Klebsiella pneumoniae* ATCC BAA 1705 and ATCC BAA 1706 were used as standard strains, and *Escherichia coli* SK191 and BL21 were used as control strains. The color in the heatmap represents the EOP. Bar charts on the X- and Y-axes represent the number of infections in each phage and host, respectively.

We observed that high EOP values indicated the formation of large plaques. Thus, we performed correlation analysis between EOP values and plaque size of eight representative phages that showed different host range pattern. Pearson’s correlation coefficients (R values) were 0.75 for øKp_1, 0.86 for øKp_7-1, 0.70 for 7-2, 0.55 for øKp_14, 0.25 for øKp_21, 0.80 for øKp_24, 0.83 for øKp_27, and 0.65 for øKp_31 (Fig. S2). Therefore, plaque size was positively correlated with the EOP.

### 3. OD_600_ kinetics of *K. pneumoniae* challenged with phages

OD_600_ kinetics were analyzed for all phage–indicator host combinations. Individual phages were added to cultures of indicator hosts at 10^8^ pfu/ml. The OD_600_ decreased 1 h after each phage was added, while that of the Ec1 and øEc_1 mixture did not decrease and approximated that of the host strain without any phages (Fig. 3). The OD_600_ in all phage combinations increased again 6–10 h after the addition of each phage. This result indicates that phage-resistant bacteria emerged in almost all the phage–host combinations.

**Fig. 3.**
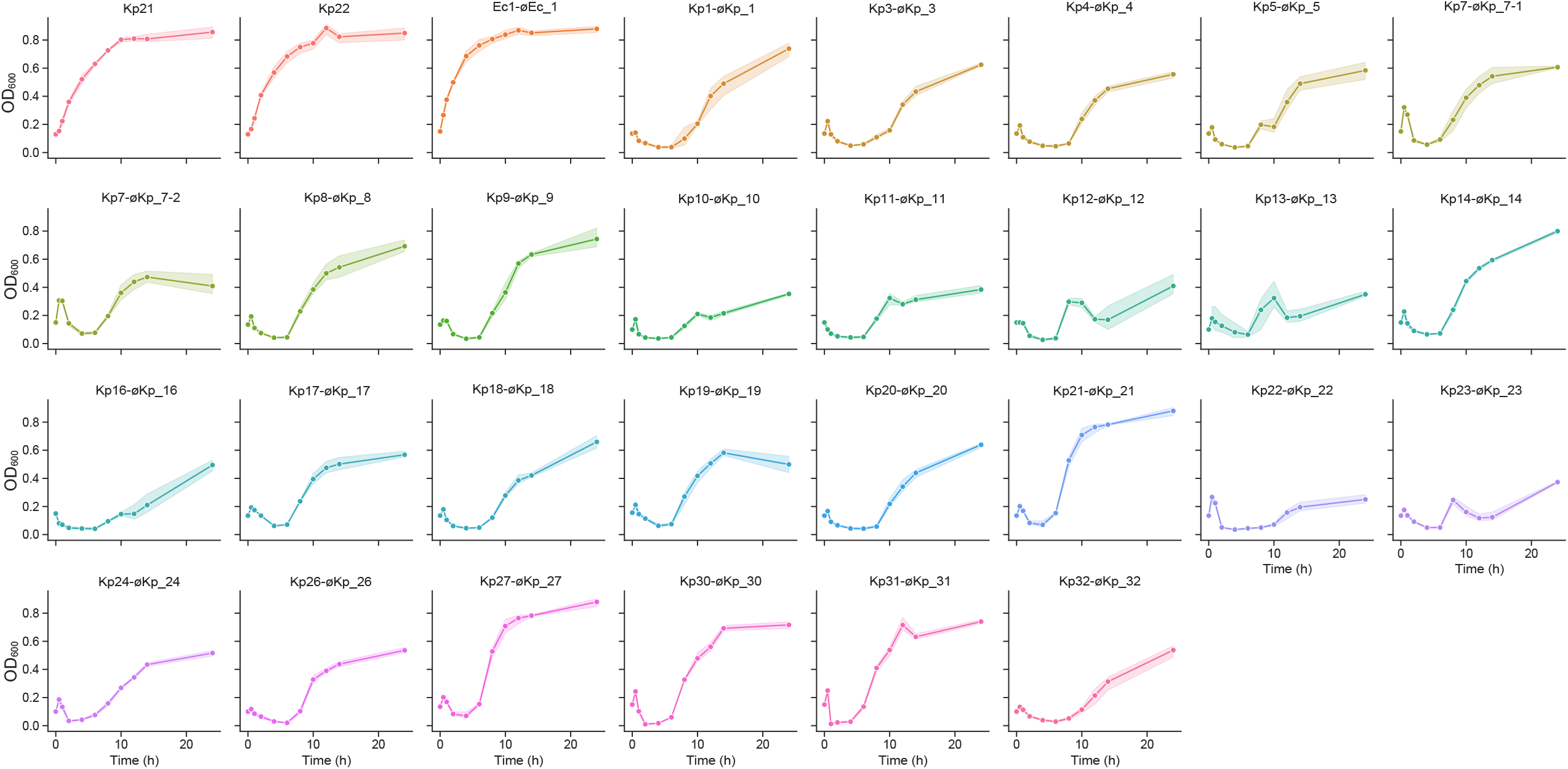
OD_600_ kinetics indicator bacteria incubated with phage. Bacterial strains were incubated up to OD_600_ = 0.1, following which each phage was added at 10^9^ pfu/ml. OD_600_ was monitored at appropriate times until 24 h. No phage was added in Kp21 and Kp22 for the negative control. All experiments in this section were performed in triplicate.

### 4. Cocktail analysis of phage-resistant bacteria Kp21

Kp21 can only be infected by øKp_21, while Kp22 can be infected by 23 phages (Fig. 2). Furthermore, øKp_21 infects 18 hosts including Kp21, but øKp_22 infects Kp22 and Kp31 strains only (Fig. 2). We focused on these contrasting strains, øKp_21 and øKp_22, for the phage cocktail experiment. The phage cocktail consisted of 10 phages (øKp_16–26), which include 8 *Tevenviridae*, 1 *Alcyoneusvirus*, and 1 *Siphoviridae*. In the Kp21 and øKp_21 combination, the OD_600_ increased again after 6 h and reached that of the negative control after 24 h (Fig. 4A). However, the OD_600_ did not increase until 14 h when Kp21 was combined with the phage cocktail (Fig. 4A). Therefore, the phage cocktail delayed the emergence of phage-resistant Kp21 in the *in vitro* assay. However, the OD_600_ of the cocktail against Kp22 increased again after 10 h, which is the same time taken as the OD_600_ of single øKp_22 (Fig. 4C). This result indicates that the phage cocktail failed to delay the emergence of phage-resistant Kp22. We isolated øKp_21-resistant Kp21 (Kp21r) and øKp_22-resistant Kp22 (Kp22r) according to the method described in the Materials and methods section. Although the OD_600_ kinetics in the mixture of Kp21r and øKp_21 did not decrease and approximated that of Kp21r without any phages, the OD_600_ reduced in the Kp21r–phage cocktail combination (Fig. 4B). This result indicates that the Kp21r strain is resistant to phage 21 but susceptible to the phage cocktail. However, OD_600_ did not decrease in either Kp22r-øKp_22 or Kp22r-cocktail combinations, suggesting that the phage-resistant Kp22r is not susceptible to either øKp_22 or the phage cocktail (Fig. 4D).

**Fig. 4.**
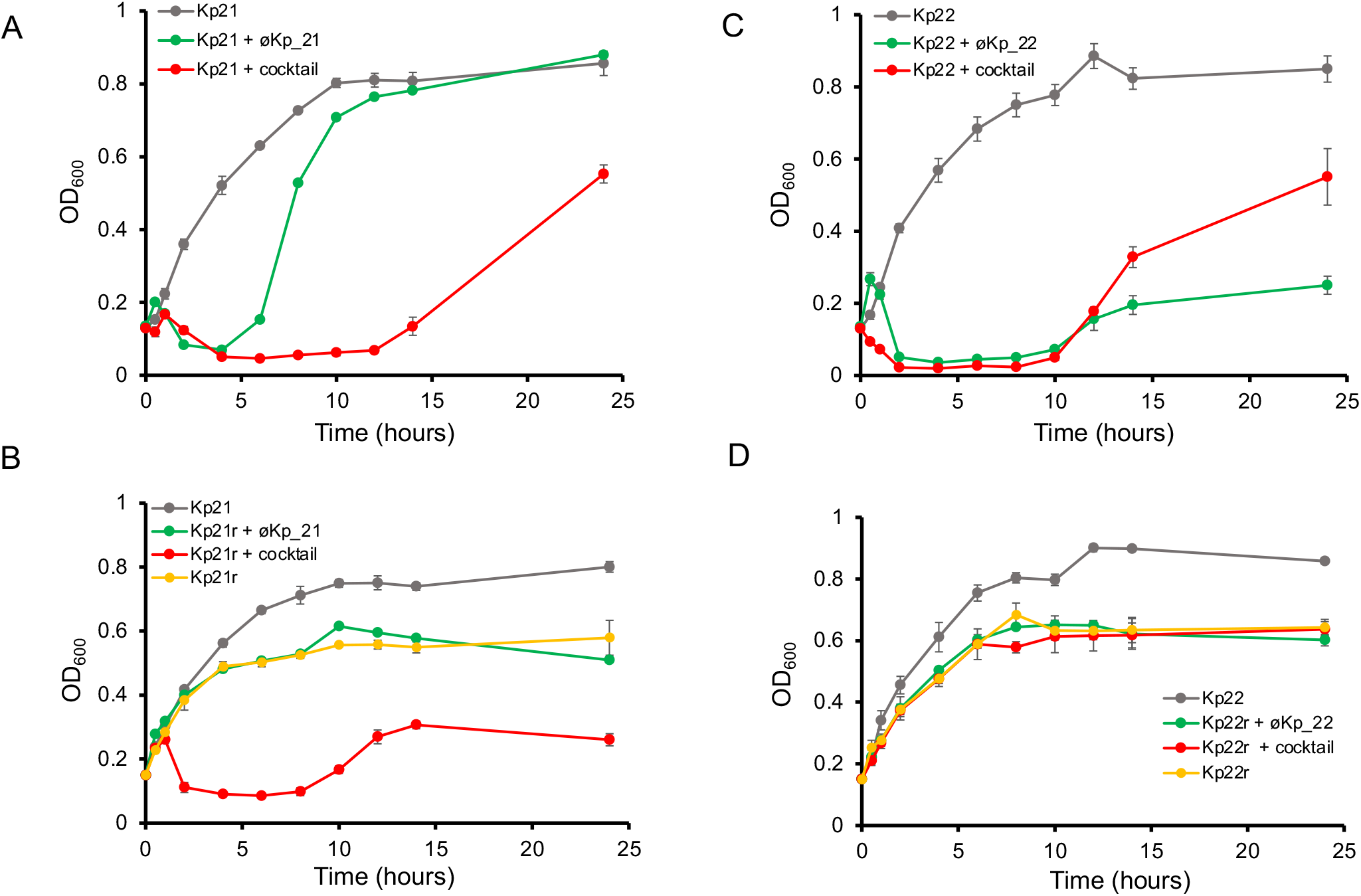
Cocktail experiment of Kp21r and Kp22r. Phage-resistant Kp21 (Kp21r) and Kp22 (Kp22r) were derived from the culture medium after 24 h incubation with øKp_21 or øKp_22. The cocktail consisted of 10 phages, and each phage was at 10^7^ pfu/ml. OD_600_ was monitored at appropriate times until 24 h. All experiments in this section were performed in triplicate.

### 5. Shifting of phage sensitivity between Kp21 and Kp21r

To analyze the susceptibility of Kp21r to the phage cocktail, we compared phage plaque formation between Kp21 and Kp21r. According to the host range analysis, no phages formed plaques on Kp21. However, the Kp21r strain became susceptible to øKp_16, 17, 18, 19, 20, 23, and 26, which are of the genus *Tevenviridae*. This result suggests that Kp21r becomes prone to be infected from other phages in compensation for resistance to øKp21 (Fig. 5A). Furthermore, the Kp21r strain showed a sparse background on LB agar plates containing *Tevenviridae* viruses (Fig. 5A). This sparse background is not caused by confluent plaque lysis. This implies a mechanism by which the phage cocktail prevents the emergence of phage-resistant bacteria without phage infection. To determine the efficiency with which phages can kill the Kp21r strain, we measured the colony-forming unit (cfu) values of Kp21 and Kp21r by mixing the individual phages used in the phage cocktail. In øKp_21, the cfu of Kp21 was reduced (10^4^ cfu/ml), but that of Kp21r was almost the same as that of the control Kp21r (10^7^–10^8^ cfu/ml). In øKp_22 and øKp_24, cfu/ml in Kp21r was not significantly decreased compared to that in Kp21. These results are consistent with our observation that these phages are incapable of infecting Kp21 and Kp21r. In other phages, the cfu of Kp21r strain significantly decreased compared to that of Kp21. In particular, colonies were not detected for øKp_18, 19, 20, or 23-Kp21r combinations. Our results indicate that Kp21r viable cells were wiped out by phages that newly infected Kp21r.

**Fig. 5.**
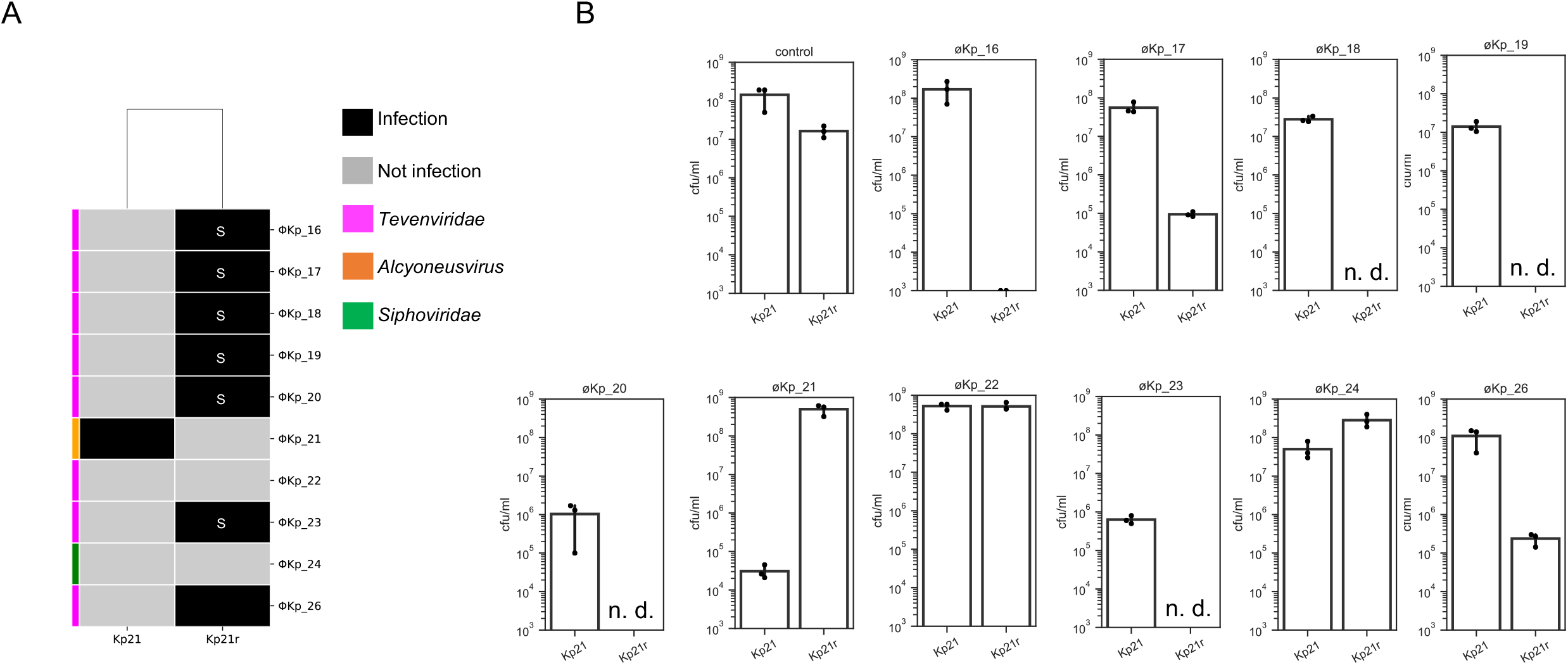
Analysis of the shifting in susceptibility to the phage cocktail in f22 and f22r. (A) the host range in f22 and f22r was investigated against 10 phages comprising a phage cocktail. “S” represents a sparse bacterial lawn. (B) Colony forming units are mentioned under each phage. f22 or f22r were mixed with individual phages, and 2 h after phage addition, samples were diluted to 10^−2^ and 10^−4^, and lawned onto LB plates. n.d. means that colonies were not detected at the 10^−2^ condition.

### 6. Characterization of Kp21 and Kp21r

To analyze the differences between the strains Kp21 and Kp21r, we performed an adsorption assay of øKp_21 for Kp21 and Kp21r. The assay showed that unadsorbed øKp_21 for the Kp21r strain was approximately 100 % at 5 min after the phage was added to the Kp21 strain, while for Kp21, the value was 2–5 % (Fig. 6A). In addition, no plaques were detected when øKp_21 was input at 10^9^ pfu (EOP < 10^−9^) (Fig. 6B). These results indicate that øKp_21 loses its ability to adsorb Kp21r. Next, we detected single nucleotide polymorphism (SNPs) between Kp21 and Kp21r. Insertion mutations were detected in two genes, *thpA* (encoding the inner membrane protein) and *cpsA* (encoding exopolysaccharide synthesis genes) as shown in Table S4; this result suggests that the øKp_21 phage recognizes the capsular polysaccharide of Kp21. Insertion mutations of *cpsA* in Kp21r occur at the 452^nd^ nucleotide position and a stop codon appeared at the 463^rd^ nucleotide position (Fig. 6C). Therefore, 141 amino acids are truncated at C-terminus CpsA of Kp21r (154 amino acids long); the wild-type CpsA amino acid length is 295. This severe truncation can result in deficient capsular polysaccharide biosynthesis of Kp21r.

**Fig. 6.**
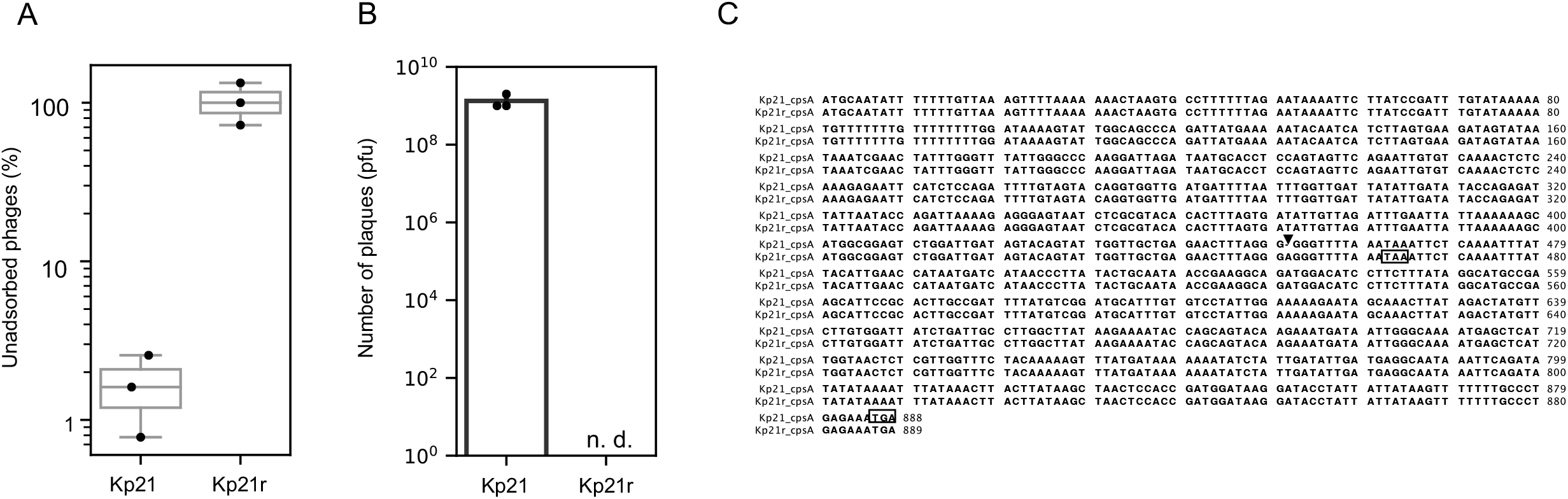
Characterization of phage-resistant Kp21r strain. (A) Adsorption assay of øKp_21 against Kp21 and Kp21r strains. Kp21 and Kp21r were incubated until OD_600_ = 0.5, following which øKp_21 was added at 10^7^ pfu/ml and incubated at 37°C with shaking at 200 rpm. After 5 min, 200 µl of the mixture was withdrawn and centrifuged at 9,100 *×g* for 1 min. The number of phages in the supernatant was measured. (B) 1.0 × 10^9^ pfu of øKp_21 was mixed with 100 µl of overnight Kp21 or Kp21r culture. Thereafter, 5 ml of 0.6 % soft agar was added to the host and phage mixture and incubated at 37°C overnight. n.d. means that plaques were not detected. (C) Nucleotide sequences of *cpsA* in Kp21 and Kp21r were aligned using ClustalW (http://clustalw.ddbj.nig.ac.jp/). Insertion mutation (A) is shown by an arrow, and the stop codons of *cspA* in f22 and f22r are shown by a square.

## Discussion

Phage therapy is increasingly being recognized as an effective strategy to combat antimicrobial-resistant bacteria, especially nosocomial pathogens (8–10, 19). A recent study demonstrated that inflammation in a mouse model of inflammatory bowel disease (IBD) was suppressed by the eradication of *K. pneumoniae* using a phage cocktail. This suggests that phage therapy targeting *K. pneumoniae* successfully treats IBD in humans (20).

In this study, we isolated and characterized novel bacteriophages targeting antimicrobial-resistant *K. pneumoniae* harboring the pKPI-6 plasmid, which encodes *bla*_IMP-6_. We isolated 29 novel phages from sewage in west Japan against the *K. pneumoniae* and *E. coli* harboring pKPI-6 plasmid. Genome sequence analysis suggests that *Tevenviridae* members in this study except øKp_22 are very similar in genome size and identity; thus, these phages were considered to be variants of the same species (Table S1). We were concerned that our phages encode antimicrobial resistance (AMR) and virulence factor (VF) genes, because phages can transfer these genes in clinical settings (21–23). However, our analysis indicated that neither AMR nor VF genes were detected in isolated phage genomes, thereby allowing the application of these phages in clinical settings.

Our host range experiment analysis showed that most of the phages, which were classified as *Myoviridae*, formed plaques on the 25 *K. pneumoniae* strains. Specific host strains (*K. pneumoniae* Kp12 to Kp20) showed higher EOP (Fig. 3) against most phages, suggesting that several strains exhibit high susceptibility to novel phages isolated from west Japan. Moreover, host range experiment results indicated that EOP and plaque size are positively correlated in several phages such as øKp_1, 7, 7-1, 14, 24, 27, and 31. To the best of our knowledge, few reports have described the correlation of these factors (24); however, we hypothesize that phages that show a larger plaque size have a greater burst size and/or adsorption efficacy. These results can facilitate the development of novel phages that have a higher virulence to the host bacteria; moreover, our work can also guide the selection of phage strains for developing a phage cocktail (25, 26).

The phage cocktail experiment revealed that Kp21r became newly susceptible to other phages in the phage cocktail. Phage cocktail analysis showed that Kp22 did not retard the emergence of phage-resistant Kp22 (Kp22r), which was in contrast to the results for the Kp21 strain. This antithetical result of the Kp21 and Kp22 cocktail experiment implies that the phage cocktail is not an all-round strategy; however, it remains the most reliable strategy to combat bacteriophages, thus far. No universal methods or guidelines have been established for developing the cocktail, and it is difficult to predict the combination of phages that can inhibit the emergence of phage-resistant bacteria with the greatest efficiency. However, it has been reported that several phages that recognize different receptors of the host should be mixed to efficiently decrease the occurrence of phage-resistant bacteria (27–29).

Our experimental results indicate that Kp21r became infected by øKp_16, 17, 18, 19, 20, 23, and 26, while Kp21 was only subject to infection by øKp_21 (Fig. 5). This result suggests that phage–resistant bacteria is easily attacked by other phages that exist in environment during evolutionary arms race. Adsorption assays explained that øKp_21 lacks the ability to adsorb Kp21r. SNP analysis of Kp21 and Kp21r revealed that insertion mutations occurred in at least two genes; *cpsA*, encoding putative capsular biosynthesis protein and *thpA*, encoding sugar ABC transporter substrate-binding protein. It has been reported that capsular polysaccharide function as the barrier to infect phages (30). A recent article reports a novel phage that recognizes the capsular polysaccharide of *K. pneumoniae* (31), and accordingly, we speculate that capsular polysaccharide is one of the factors allowing øKp_21 to adsorb to its host Kp21. We found that *Myoviridae*, in the phage cocktail, diminished the lawn density of Kp21r on the plates. We posit that this phenomenon was caused by lysis mechanisms such as “lysis from without (LO)” or “rapid lysis” (32, 33). Gp5 in T4 phage, which encodes tail lysozyme, is known to cause LO (34). Gp5 forms the complex with T4 phage tail and when T4 phage adsorbs to their host, Gp5 degrades the peptidoglycan layer. We found that members of *Myoviridae*, used for the phage cocktail in this study, encode the baseplate with tail lysozyme (Table S3), which has the capability of peptidoglycan degradation. SNPs analysis of Kp21 and Kp21r suggests that Kp21r possesses deficient capsular polysaccharide, and thus, phage tail protein encoded in *Myoviridae* may more efficiently degrade Kp21r peptidoglycan and result in rapid lysis. Some studies have reported that phage-resistant bacteria become susceptible to antibiotic due to mutation in the genes involved in antibiotics resistance and showed that phages and antibiotics combination effectively kills the target bacteria (35–38). Our results demonstrate that the combinations of phage and phage-encoded tail lysozyme efficiently eliminate and/or inhibit the phage-resistant bacterial growth (39).

In conclusion, we isolated and characterized novel phages infecting *K. pneumoniae* and *E. coli* harboring the pKPI-6 plasmid; this is the first step in the construction of a public phage bank and phage therapy in Japan. Overall, our phage sets can diminish the threat of *K. pneumoniae* harboring the pKPI-6 plasmid isolated in clinical settings. Our phage sets also contain an adequate number of phage types for developing a phage cocktail. However, the development of a high-throughput method is required for efficiently isolating additional novel phages.

## Materials and Methods

### 1. Phage isolation and host information

Thirty-two IMP-6-producing isolates of *K. pneumoniae* and one IMP-6-producing isolate of *E. coli* were isolated from clinical settings in western Japan. Further, 29 novel phages were isolated from sewage in west Japan. Briefly, 100 µl of sewage was mixed with an overnight culture of the indicator host, and the mixture added to 3 ml of YT-soft agar prior to inoculation onto Luria–Bertani (LB) agar. The plates were incubated at 37°C overnight, and thereafter, single-plaque isolation was performed. Plaques were suspended in 1 ml LB medium and incubated for 2 h. Next, 50 µl chloroform (Fujifilm Wako Pure Chemical Corporation, Osaka, Japan) was added to each solution; the mixture was vortexed and then centrifuged at 10,000 *×g* for 10 min at 4°C. Supernatants and individual indicator hosts were mixed and incubated at 37°C overnight on LB agar plates. The single-plaque isolation procedure was repeated three times, and isolated phages were stored at 4°C until use. Kp21 was renamed from the *K. pneumoniae* f22 strain.

### 2. Phage propagation and purification

Pre-cultured host strains were inoculated into 3 ml fresh LB medium (1:100) and incubated at 37°C until the OD_600_ reached 0.5. Thereafter, each phage that was originally isolated using the indicated host was added and incubated at 37°C with shaking at 200 rpm for 4–6 h. Following lysis, 50 µl chloroform (FUJIFILM Wako Pure Chemical Corporation) was added to 1 ml of the phage lysate, vortexed, and then centrifuged at 9,100 *×g* for 10 min. Supernatants were filtered using a 0.22 µm pore-size membrane (Millipore, MA, USA). Cesium chloride (CsCl) density gradient phage purification was performed as described previously (40, 41) with some modification. Briefly, 10% polyethylene glycol 6000 (FUJIFILM Wako Pure Chemical Corporation) and 0.5 M NaCl were added to phage lysates and kept at 4°C for 1.5 h. Thereafter, phage lysates were centrifuged at 10,000 ×*g* for 30 min. Phage pellets were suspended in 1 ml TM buffer (10 mM Tris-HCl, and 5 mM MgCl_2_ [pH 7.5]), and 100 µg/ml DNase I (Roche, Basel, Switzerland) and RNase I (Thermo Fisher Scientific, MA, USA) were added to the phage solution and incubated at 37°C for 30 min. CsCl (FUJIFILM Wako Pure Chemical Corporation) at three different weights (ρ = 1.3, 1.5, and 1.7) and phage solution were overlaid in tubes and ultracentrifuged (Optima MAX-TL; Beckman Coulter, California, USA) at 100,000 *×g* for 1 h. Phage bands were then collected and dialyzed in SM buffer (25 mM Tris-HCl [pH, 7.5], 100 mM NaCl, and 8 mM MgSO_4_).

### 3. Phage propagation and electron microscopic imaging

Copper mesh grids coated with formvar and carbon (Veco grids; Nissin EM, Tokyo, Japan) were glow-discharged and placed on drops of the phages for 1 min. Thereafter, they were rinsed with distilled water and stained with a 2% uranyl acetate solution. Samples were examined using transmission electron microscopy (HT7700; Hitachi Ltd., Tokyo, Japan) at 80 kV.

### 4. Host range determination and EOP assay

Each host was incubated at 37°C overnight, and 100 µl of each overnight culture was mixed with 100 µl of each phage. Thereafter, 5 ml of 0.6 % soft agar was added to the host–phage mixture and inoculated onto LB agar. The plates were then incubated at 37°C overnight, and the number of plaques in each plate counted. EOP was calculated using the formula below:

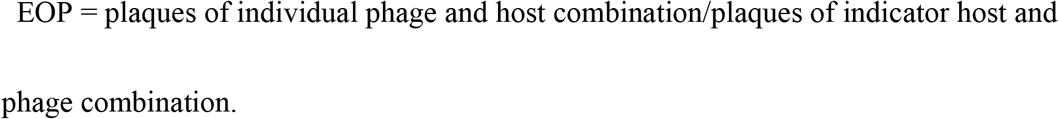

The detection limit of EOP was set as 10^−4^ pfu. EOP was measured for all phage–bacteria combinations. Each plaque image was taken using scan1200 (Interscience, Montpellier, France) for measuring plaque sizes. Plaque size (mm^2^) was measured using Fiji (https://fiji.sc) version 2.3.0, with 1 mm being 11 pixels. For very small plaques, the edge of individual plaques was detected using the “find edge” tool in Fiji. Ten plaque areas were measured in each phage–host combination if the number of plaques on the plate was more than 10.

### 5. OD_600_ kinetics and cocktail experiment

The host colony was pre-cultured in LB medium overnight at 37°C. Subsequently, the pre-cultured bacteria were inoculated (1:100) into fresh LB medium and incubated at 37°C with shaking at 200 rpm until OD_600_ = 0.1. Each indicated phage was added to the culture at 1.0 × 10^8^ pfu/ml and the mixed culture incubated at 37°C with shaking at 200 rpm. OD_600_ was measured at appropriate times for 24 h. In phage cocktail experiments, 10 phages (øKp_16, 17, 18, 19, 20, 21, 22, 23, 24, and 26) were mixed at 1.0 × 10^7^ pfu/ml of each individual phage. All experiments in this section were performed in triplicate.

### 6. Host and phage genome sequences

All phage genomic DNA was extracted using Norgen phage DNA isolation kit (Norgen Biotak, Birmingham, UK) following the manufacturer’s instructions. Each phage DNA library was constructed using the QIA seq FX DNA library kit (Qiagen), and sequencing was performed using the Illumina MiSeq platform. Genome assembly was performed using Shovill with default settings. Phage contigs were filtered with the contig length < 200 and coverage < 25. Bacteria strains and phage strains were annotated using prokka (42) or PGAP (43) version 2021-07-01.build5508. For Nanopore long-read sequencing, we used the Monarch HMW DNA Extraction Kit for Tissue (NEB, MA, USA) following the manufacturer’s instructions.

A long-read library was prepared using the Rapid Barcoding kit (Oxford Nanopore Technologies, Oxford, United Kingdom, catalog number: SQK-RBK004) and sequenced with an R9 flow cell (Oxford Nanopore Technologies, catalog number: FLO-MIN106) and a GridION device (Oxford Nanopore Technologies). Basecalling was performed using Guppy version 5.0.12 with high accuracy mode. The obtained long reads and MiSeq short reads that were trimmed using fastp v0.20.1 were assembled using Unicycler v0.4.8 with default parameters. Annotation was conducted using PGAP version 2021-07-01.build5508.

### 7. Bioinformatics analysis

For protein prediction in phages, we constructed the phage protein databases from International Committee on Taxonomy of Viruses (ICTV), which consists of 4,312 genomes and 4,62,579 proteins (https://www.ncbi.nlm.nih.gov/genomes/GenomesGroup.cgi?taxid=28883) (September 2021) using local blastp (44) with the e-value threshold < 2e-20. Protein domains were detected using hmmer (https://www.ebi.ac.uk/Tools/hmmer/) with the Pfam-A 35.0 database at e-value of < 1e-10. Phage classification was determined according to National Center for Biotechnology Information (NCBI) GenBank and ICTV. Average nucleotide identity was conducted using the average_nucleotide_identity.py program in pyani packages (45). MUMmer was used to align nucleotide sequences. AMR genes and virulence genes were detected using ABRicate version 1.0.1 (https://github.com/tseemann/abricate) under default settings. The ResFinder database was used to extract AMR genes (46), and the Virulence Factors Database (VFDB) was used to extract virulence genes (47). The packaging mechanism and terminal repeats were analyzed using Phagetermvirome version 4.0.1 (48), and tRNA was detected using tRNAscan-SE 2.0. (49). Kp21 and Kp21r SNPs analysis was performed using SNIPPY at default settings (50).

### 8. Phage-resistant Kp21 (Kp21r) and Kp22 (Kp22r) strain

Kp21 and Kp22 were cultured with øKp_21or øKp_22, respectively at 37°C. After 24 h of incubation, 1 ml of each culture was centrifuged at 4,400 *×g* for 10 min. The supernatant was discarded and the pellets washed with LB medium. This procedure was repeated twice. Thereafter, the pellet was resuspended using saline (0.85 % NaCl) and the suspension plated on LB agar and incubated at 37°C overnight. A single colony was incubated overnight at 37°C with shaking at 200 rpm. Glycerol stock of Kp21r culture was stored at – 80°C until use.

### 9. Adsorption assay

Kp21 and Kp21r were cultured in LB medium and incubated at 37°C until OD_600_ = 0.5. Subsequently, 3.0 × 10^6^ pfu øKp21 was added and incubated at 37°C with shaking at 200 rpm for 5 min. Next, 20 µl chloroform was added to 200 µl of the mixture and vortexed. The samples were then centrifuged at 9,100 *×g* for 1 min and the supernatant collected. A 100 µl aliquot of the supernatant was mixed with Kp21 and plaque assays performed to measure the number of unadsorbed phages. The percentage of the unadsorbed phages was calculated using the formula below:

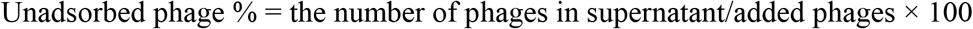

### 10. Characterization of switched phage sensitivity between Kp21 and Kp21r

The phage sensitivity of Kp21 and Kp21r were examined. Briefly, 1.0 × 10^9^ pfu of individual phages used in the phage cocktail was mixed with 100 µl of overnight Kp21 or Kp21r culture. Thereafter, 5 ml of 0.6 % soft agar was added to the host and phage mixture and then poured onto LB agar and incubated at 37°C overnight. The colony count was examined as follows. Kp21 and Kp21r were incubated at 37°C until OD_600_ = 0.1. Subsequently, Kp21 was added to 1.0 × 10^9^ pfu, and the mixture was incubated at 37°C. The mixture was collected 2 h after phages were added and centrifuged at 3,300 *×g* for 15 min. The supernatant was discarded, and the pellet was suspended in 500 µl phosphate-buffered saline (0.137 M NaCl, 0.27 mM KCl, 0.1 M Na_2_HPO_4_, 18 mM KH_2_PO_4_). The suspension was diluted to 10^−2^ and 10^−4^, and 100 µl of these dilutions was lawned onto LB agar.

## 11. Data availability

Row sequence reads for all phages were deposited to DDBJ/EMBL/GenBank under Bioproject (number: PRJDB14376), and DRA numbers are listed in Table S1. Complete genome sequences of Kp21 and Kp21r were deposited in GenBank (accession numbers AP026912 and AP026913).

## Acknowledgment

This work was partially supported by JSPS KAKENHI (21K1633 to K.K., 18K08455 and 22K08592 to M.K.) from the Ministry of Education, Culture, Sports, Science and Technology (MEXT), Japan; grants (19lm0203008j0003 A058 and 22ym0126811j0001 A255 to M.K.) from the Japan Agency for Medical Research and Development (AMED) and grants (JP21fk0108604j0001, 22fk0108604j0002, 21fk0108132j0002 to M.S.) from Research Program on Emerging and Re-emerging infection Diseases (AMED).

## Tables

Table S1 The genomic information of phages isolated in Japan. Genome assembly was performed using Shovill with default settings. Phage contigs were filtered with the contig length < 200 and/or coverage < 25. “CDS after filtered” and “contigd_filtered” columns represent the number of CDS from the filtered contigs. The packaging mechanism and terminal repeats in each phage were presumed using Phagetermvirome.

Table S2 The table of host range and EOP for all host-phage combination. EOP was calculated by plaques of individual phage and host combination divided by plaques of indicator host and phage combination.

Table S3 The list of the domain prediction for each phage protein. Pfam-A 27 was used as phage database.

Table S4 The list of SNPs between Kp21 and Kp21r. SNPs were detected using Snippy as default settings.

---

**Fig. S1.**
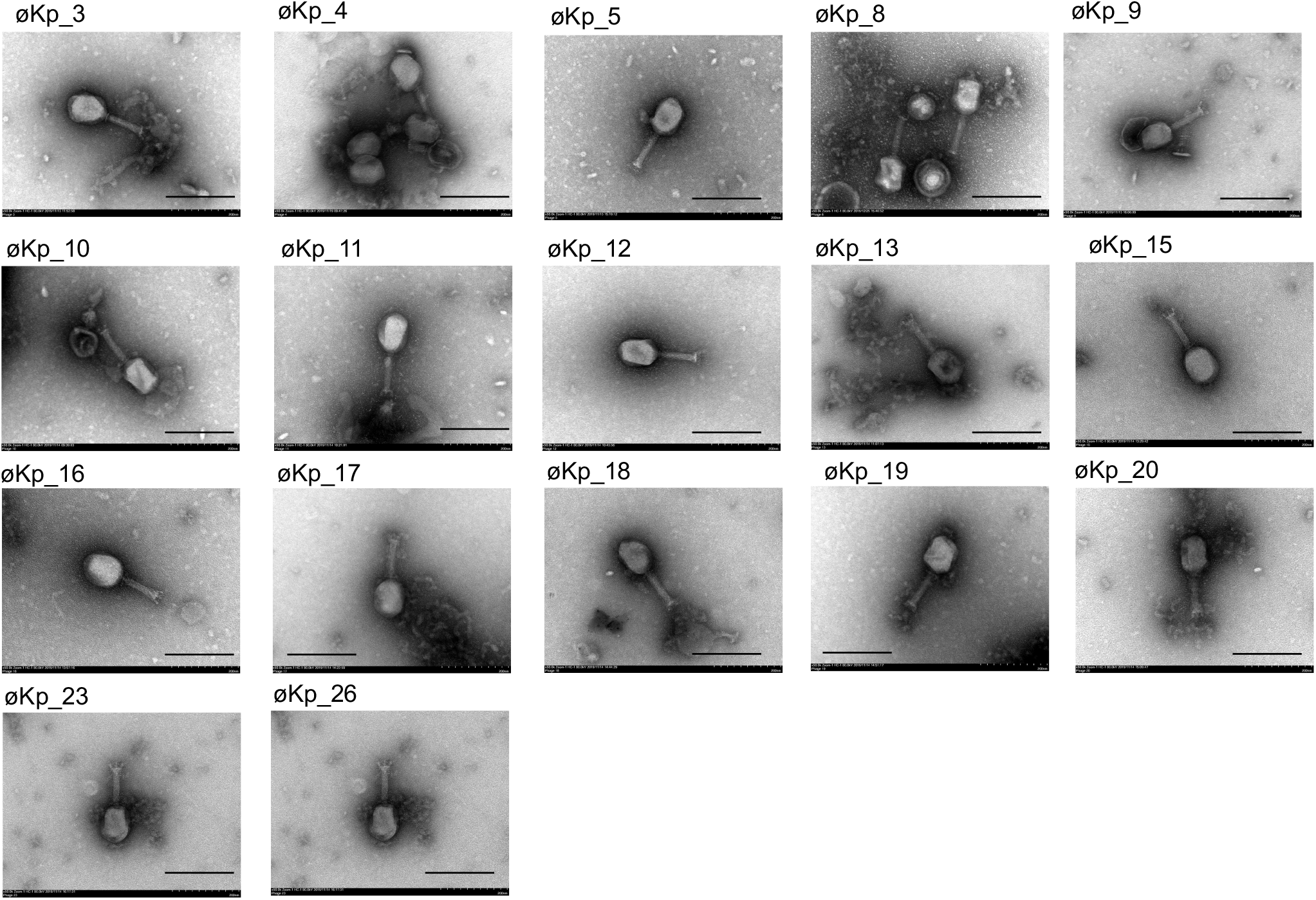
All TEM images of *Myoviridae* in this study. The bar represents 200 nm in individual images.

**Fig. S2.**
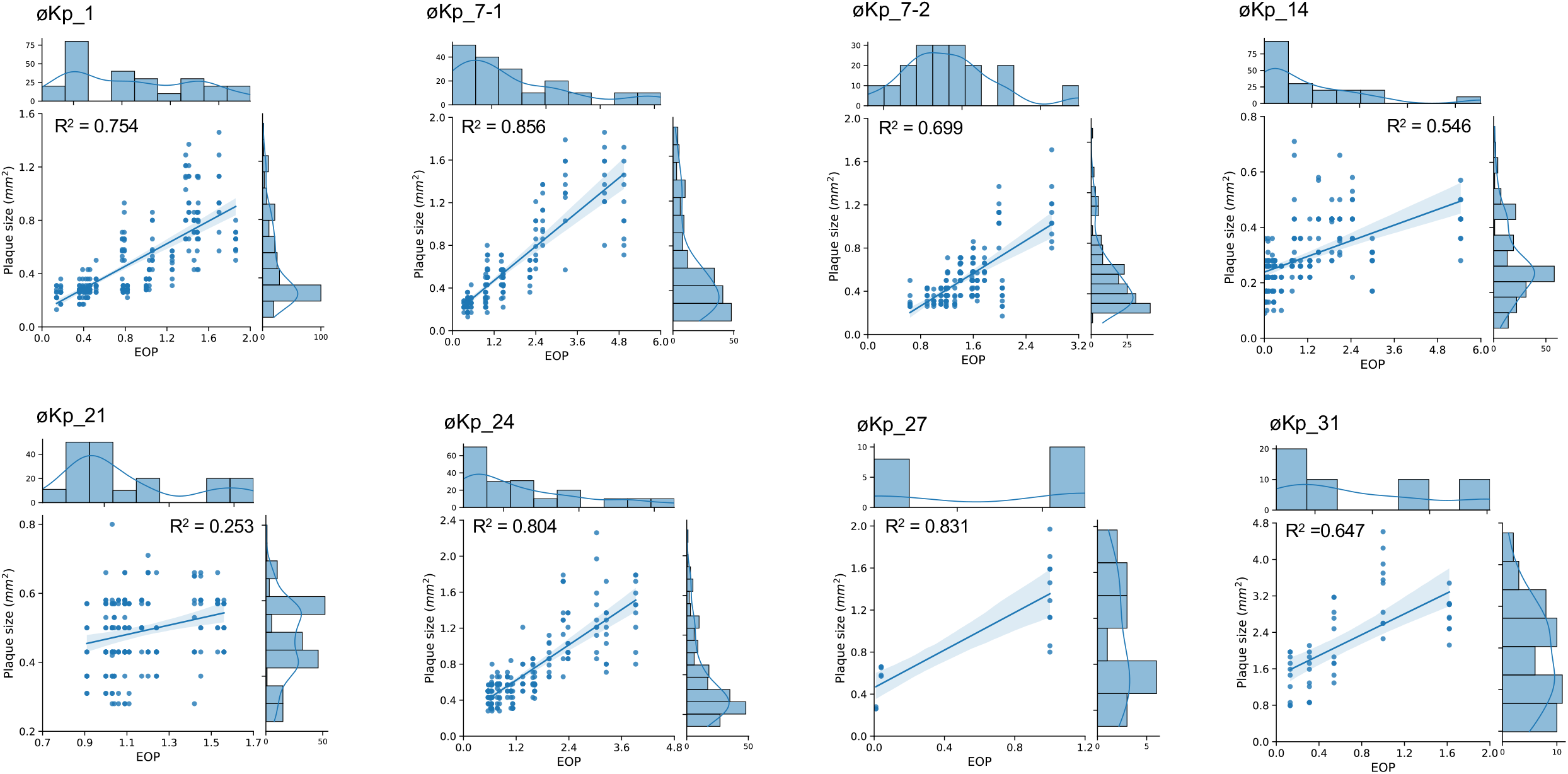
Correlation between plaque size and EOP. Plaque size (mm^2^) and EOP in the phages were measured in representative phage strains. The X- and Y-axes show the EOP in each phage and plaque size, respectively. A maximum of 10 plaques were randomly selected in individual phage-host combinations and plaque sizes were measured using ImageJ. Python packaging Seaborn was used to visualize the correlation, and R correlation in each combination was calculated using Scipy version 1. 8. 1.

## References

1. Effah CY, Sun T, Liu S, Wu Y. 2020. Klebsiella pneumoniae: An increasing threat to public health. Ann Clin Microbiol Antimicrob 19:1–9.

2. Yamagishi T, Matsui M, Sekizuka T, Ito H, Fukusumi M, Uehira T, Tsubokura M, Ogawa Y, Miyamoto A, Nakamori S, Tawa A, Yoshimura T, Yoshida H, Hirokawa H, Suzuki S, Matsui T, Shibayama K, Kuroda M, Oishi K. 2020. A prolonged multispecies outbreak of IMP-6 carbapenemase-producing Enterobacterales due to horizontal transmission of the IncN plasmid. Sci Rep 10:1–9.

3. Abe R, Akeda Y, Sugawara Y, Takeuchi D, Matsumoto Y, Motooka D, Yamamoto N, Kawahara R, Tomono K, Fujino Y, Hamada S. 2020. Characterization of the Plasmidome Encoding Carbapenemase and Mechanisms for Dissemination of Carbapenem-Resistant Enterobacteriaceae. mSystems 5.

4. Nordmann P, Naas T, Poirel L. 2011. Global spread of carbapenemase producing Enterobacteriaceae. Emerg Infect Dis 17:1791–1798.

5. Kayama S, Shigemoto N, Kuwahara R, Oshima K, Hirakawa H, Hisatsune J, Jové T, Nishio H, Yamasaki K, Wada Y, Ueshimo T, Miura T, Sueda T, Onodera M, Yokozaki M, Hattori M, Ohge H, Sugai M. 2015. Complete nucleotide sequence of the IncN plasmid encoding imp-6 and CTX-M-2 from emerging carbapenem-resistant Enterobacteriaceae in japan. Antimicrob Agents Chemother 59:1356–1359.

6. Shigemoto N, Kuwahara R, Kayama S, Shimizu W, Onodera M, Yokozaki M, Hisatsune J, Kato F, Ohge H, Sugai M. 2012. Emergence in Japan of an imipenem-susceptible, meropenem-resistant Klebsiella pneumoniae carrying bla IMP-6. Diagn Microbiol Infect Dis 72:109–112.

7. Cao F, Wang X, Wang L, Li Z, Che J, Wang L, Li X, Cao Z, Zhang J, Jin L, Xu Y. 2015. Evaluation of the efficacy of a bacteriophage in the treatment of pneumonia induced by multidrug resistance klebsiella pneumoniae in mice. Biomed Res Int 2015.

8. Petrovic Fabijan A, Lin RCY, Ho J, Maddocks S, Ben Zakour NL, Iredell JR, Khalid A, Venturini C, Chard R, Morales S, Sandaradura I, Gilbey T. 2020. Safety of bacteriophage therapy in severe Staphylococcus aureus infection. Nat Microbiol 5:465–472.

9. Dedrick RM, Guerrero-Bustamante CA, Garlena RA, Russell DA, Ford K, Harris K, Gilmour KC, Soothill J, Jacobs-Sera D, Schooley RT, Hatfull GF, Spencer H. 2019. Engineered bacteriophages for treatment of a patient with a disseminated drug-resistant Mycobacterium abscessus. Nat Med 25:730–733.

10. Eskenazi A, Lood C, Wubbolts J, Hites M, Balarjishvili N, Leshkasheli L, Askilashvili L, Kvachadze L, van Noort V, Wagemans J, Jayankura M, Chanishvili N, de Boer M, Nibbering P, Kutateladze M, Lavigne R, Merabishvili M, Pirnay JP. 2022. Combination of pre-adapted bacteriophage therapy and antibiotics for treatment of fracture-related infection due to pandrug-resistant Klebsiella pneumoniae. Nat Commun 13.

11. Oechslin F. 2018. Resistance development to bacteriophages occurring during bacteriophage therapy. Viruses 10.

12. Oechslin F, Piccardi P, Mancini S, Gabard J, Moreillon P, Entenza JM, Resch G, Que YA. 2017. Synergistic interaction between phage therapy and antibiotics clears Pseudomonas Aeruginosa infection in endocarditis and reduces virulence. J Infect Dis 215:703–712.

13. Schooley RT, Biswas B, Gill JJ, Hernandez-Morales A, Lancaster J, Lessor L, Barr JJ, Reed SL, Rohwer F, Benler S, Segall AM, Taplitz R, Smith DM, Kerr K, Kumaraswamy M, Nizet V, Lin L, McCauley MD, Strathdee SA, Benson CA, Pope RK, Leroux BM, Picel AC, Mateczun AJ, Cilwa KE, Regeimbal JM, Estrella LA, Wolfe DM, Henry MS, Quinones J, Salka S, Bishop-Lilly KA, Young R, Hamilton T. 2017. Development and use of personalized bacteriophage-based therapeutic cocktails to treat a patient with a disseminated resistant Acinetobacter baumannii infection. Antimicrob Agents Chemother 61.

14. Yerushalmy O, Khalifa L, Gold N, Rakov C, Alkalay-Oren S, Adler K, Ben-Porat S, Kraitman R, Gronovich N, Ginat KS, Abdalrhman M, Coppenhagen-Glazer S, Nir-Paz R, Hazan R. 2020. The israeli phage bank (IPB). Antibiotics 9:1–7.

15. Nagel T, Musila L, Muthoni M, Nikolich M, Nakavuma JL, Clokie MR. 2022. Phage banks as potential tools to rapidly and cost-effectively manage antimicrobial resistance in the developing world. Curr Opin Virol 53:101208.

16. Savalia D, Westblade LF, Goel M, Florens L, Kemp P, Akulenko N, Pavlova O, Padovan JC, Chait BT, Washburn MP, Ackermann HW, Mushegian A, Gabisonia T, Molineux I, Severinov K. 2008. Genomic and Proteomic Analysis of phiEco32, a Novel Escherichia coli Bacteriophage. J Mol Biol 377:774–789.

17. Mirzaei MK, Eriksson H, Kasuga K, Haggård-Ljungquist E, Nilsson AS. 2014. Genomic, proteomic, morphological, and phylogenetic analyses of vB-EcoP-SU10, a Podoviridae phage with C3 morphology. PLoS One 9:1–19.

18. Koonjan S, Cooper CJ, Nilsson AS. 2021. Complete genome sequence of vb_ecop_su7, a Podoviridae coliphage with the rare c3 morphotype. Microorganisms 9:1–11.

19. Terwilliger A, Clark J, Karris M, Hernandez-Santos H, Green S, Aslam S, Maresso A. 2021. Phage therapy related microbial succession associated with successful clinical outcome for a recurrent urinary tract infection. Viruses 13.

20. Federici S, Kredo-Russo S, Valdés-Mas R, Kviatcovsky D, Weinstock E, Matiuhin Y, Silberberg Y, Atarashi K, Furuichi M, Oka A, Liu B, Fibelman M, Weiner IN, Khabra E, Cullin N, Ben-Yishai N, Inbar D, Ben-David H, Nicenboim J, Kowalsman N, Lieb W, Kario E, Cohen T, Geffen YF, Zelcbuch L, Cohen A, Rappo U, Gahali-Sass I, Golembo M, Lev V, Dori-Bachash M, Shapiro H, Moresi C,Cuevas-Sierra A, Mohapatra G, Kern L, Zheng D, Nobs SP, Suez J, Stettner N, Harmelin A, Zak N,Puttagunta S, Bassan M, Honda K, Sokol H, Bang C, Franke A, Schramm C, Maharshak N, Sartor RB,Sorek R, Elinav E. 2022. Targeted suppression of human IBD-associated gut microbiota commensalsby phage consortia for treatment of intestinal inflammation. Cell 185:2879-2898.e24.

21. Anand T, Bera BC, Vaid RK, Barua S, Riyesh T, Virmani N, Hussain M, Singh RK, Tripathi BN. 2016. Abundance of antibiotic resistance genes in environmental bacteriophages. J Gen Viro l97:3458–3466.

22. Gómez-Gómez C, Blanco-Picazo P, Brown-Jaque M, Quirós P, Rodríguez-Rubio L, Cerdà-Cuellar M,Muniesa M. 2019. Infectious phage particles packaging antibiotic resistance genes found in meatproducts and chicken feces. Sci Rep 9:1–11.

23. Kohei K, Mitsuoki K, Motoyuki S, Mariana C. 2021. Distribution of Antimicrobial Resistance andVirulence Genes within the Prophage-Associated Regions in Nosocomial Pathogens. mSphere 0:e00452–21.

24. Haines MEK, Hodges FE, Nale JY, Mahony J, van Sinderen D, Kaczorowska J, Alrashid B, Akter M,Brown N, Sauvageau D, Sicheritz-Pontén T, Thanki AM, Millard AD, Galyov EE, Clokie MRJ. 2021. Analysis of Selection Methods to Develop Novel Phage Therapy Cocktails Against AntimicrobialResistant Clinical Isolates of Bacteria. Front Microbiol 12:1–15.

25. Molina F, Simancas A, Ramírez M, Tabla R, Roa I, Rebollo JE. 2021. A New Pipeline for DesigningPhage Cocktails Based on Phage-Bacteria Infection Networks. Front Microbiol 12:1–14.

26. Molina F, Flores MM, Fernández L, Rodríguez MAV. 2022. Systematic analysis of putative phage -phage interactions on minimum - sized phage cocktails. Sci Rep 1–12.

27. Tanji Y, Shimada T, Yoichi M, Miyanaga K, Hori K, Unno H. 2004. Toward rational control ofEscherichia coli O157:H7 by a phage cocktail. Appl Microbiol Biotechnol 64:270–274.

28. Hesse S, Rajaure M, Wall E, Johnson J, Bliskovsky V, Gottesman S. 2020. Phage Resistance inMultidrug-Resistant Klebsiella pneumoniae ST258 Evolves via Diverse Mutations That Culminate inImpaired Adsorption. MBio 11:1–14.

29. Gordillo Altamirano FL, Barr JJ. 2021. Unlocking the next generation of phage therapy: the key is inthe receptors. Curr Opin Biotechnol 68:115–123.

30. Olszak T, Shneider MM, Latka A, Maciejewska B, Browning C, Sycheva L V., Cornelissen A, Danis-Wlodarczyk K, Senchenkova SN, Shashkov AS, Gula G, Arabski M, Wasik S, Miroshnikov KA,Lavigne R, Leiman PG, Knirel YA, Drulis-Kawa Z. 2017. The O-specific polysaccharide lyase fromthe phage LKA1 tailspike reduces Pseudomonas virulence. Sci Rep 7:1–14.

31. Hao G, Shu R, Ding L, Chen X, Miao Y, Wu J, Zhou H, Wang H. 2021. Bacteriophage SRD2021recognizing capsular polysaccharide shows therapeutic potential in serotype K47 Klebsiellapneumoniae infections. Antibiotics 10.

32. Chegini Z, Khoshbayan A, Taati Moghadam M, Farahani I, Jazireian P, Shariati A. 2020. Bacteriophage therapy against Pseudomonas aeruginosa biofilms: A review. Ann Clin MicrobiolAntimicrob 19:1–17.

33. Liao Y-T, Zhang Y, Salvador A, Harden LA, Wu VCH. 2022. Characterization of a T4-likeBacteriophage vB_EcoM-Sa45lw as a Potential Biocontrol Agent for Shiga Toxin-ProducingEscherichia coli O45 Contaminated on Mung Bean Seeds. Microbiol Spectr 10.

34. Ye N, Nemoto N. 2004. Processing of the tail lysozyme (gp5) of bacteriophage T4. J Bacteriol 186:6335–6339.

35. Burmeister AR, Fortier A, Roush C, Lessing AJ, Bender RG, Barahman R, Grant R, Chan BK, TurnerPE. 2020. Pleiotropy complicates a trade-off between phage resistance and antibiotic resistance. ProcNatl Acad Sci U S A 117.

36. Gurney J, Pradier L, Griffin JS, Gougat-Barbera C, Chan BK, Turner PE, Kaltz O, Hochberg ME. 2020. Phage steering of antibiotic-resistance evolution in the bacterial pathogen, Pseudomonasaeruginosa. Evol Med Public Heal 2020:148–157.

37. Liu CG, Green SI, Min L, Clark JR, Salazar KC, Terwilliger AL, Kaplan HB, Trautner BW, RamigRF, Maresso AW. 2020. Phage-antibiotic synergy is driven by a unique combination of antibacterialmechanism of action and stoichiometry. MBio 11:1–19.

38. Nakamura K, Fujiki J, Nakamura T, Furusawa T, Gondaira S, Usui M, Higuchi H, Tamura Y, Iwano H. 2021. Fluctuating Bacteriophage-induced galU Deficiency Region is Involved in Trade-off Effectson the Phage and Fluoroquinolone Sensitivity in Pseudomonas aeruginosa. Virus Res 306:198596.

39. Roach DR, Donovan DM. 2015. Antimicrobial bacteriophage-derived proteins and therapeuticapplications. Bacteriophage 5:e1062590.

40. Uchiyama J, Rashel M, Maeda Y, Takemura I, Sugihara S, Akechi K, Muraoka A, Wakiguchi H,Matsuzaki S. 2008. Isolation and characterization of a novel Enterococcus faecalis bacteriophageφEF24C as a therapeutic candidate. FEMS Microbiol Lett 278:200–206.

41. Kitamura N, Sasabe E, Matsuzaki S, Daibata M, Yamamoto T. 2020. Characterization of two newlyisolated Staphylococcus aureus bacteriophages from Japan belonging to the genus Silviavirus. ArchVirol 165:2355–2359.

42. Seemann T. 2014. Prokka: Rapid prokaryotic genome annotation. Bioinformatics 30:2068–2069.

43. Tatusova T, Dicuccio M, Badretdin A, Chetvernin V, Nawrocki P, Zaslavsky L, Lomsadze A, Pruitt KD, Borodovsky M, Ostell J. 2016. NCBI prokaryotic genome annotation pipeline 44:6614–6624.

44. Altschul SF, Gish W, Miller W, Myers EW, Lipman DJ. 1990. Basic local alignment search tool. JMol Biol 215:403–410.

45. Pritchard L, Glover RH, Humphris S, Elphinstone JG, Toth IK. 2016. Genomics and taxonomy indiagnostics for food security: Soft-rotting enterobacterial plant pathogens. Anal Methods 8:12–24.

46. Zankari E, Hasman H, Cosentino S, Vestergaard M, Rasmussen S, Lund O, Aarestrup FM, Larsen MV. 2012. Identification of acquired antimicrobial resistance genes. J Antimicrob Chemother 67:2640–2644.

47. Chen L, Zheng D, Liu B, Yang J, Jin Q. 2016. VFDB 2016: Hierarchical and refined dataset for bigdata analysis - 10 years on. Nucleic Acids Res 44:D694–D697.

48. Garneau JR, Legrand V, Marbouty M, Press MO, Vik DR, Fortier LC, Sullivan MB, Bikard D, Monot M. 2021. High-throughput identification of viral termini and packaging mechanisms in viromedatasets using PhageTermVirome. Sci Rep 11:1–9.

49. Chan PP, Lin BY, Mak AJ, Lowe TM. 2021. TRNAscan-SE 2.0: Improved detection and functionalclassification of transfer RNA genes. Nucleic Acids Res 49:9077–9096.

50. Page AJ, Taylor B, Delaney AJ, Soares J, Seemann T, Keane JA, Harris SR. 2016. SNP-sites: rapidefficient extraction of SNPs from multi-FASTA alignments. Microb genomics 2:e000056.

